# Neural decoding from surface high-density EMG signals: influence of anatomy and synchronization on the number of identified motor units

**DOI:** 10.1101/2022.02.05.479100

**Authors:** Daniela Souza de Oliveira, Andrea Casolo, Thomas G. Balshaw, Sumiaki Maeo, Marcel Bahia Lanza, Neil R.W. Martin, Nicola Maffulli, Thomas Mehari Kinfe, Björn Eskofier, Jonathan P. Folland, Dario Farina, Alessandro Del Vecchio

## Abstract

**Objective:** High-density surface electromyography (HD-sEMG) allows the reliable identification of individual motor unit (MU) action potentials. Despite the accuracy in decomposition, there is a large variability in the number of identified MUs across individuals and exerted forces. Here we present a systematic investigation of the anatomical and neural factors that determine this variability.

**Approach:** We investigated factors of influence on HD-sEMG decomposition, such as synchronization of MU discharges, distribution of MU territories, muscle-electrode distance (MED - subcutaneous fat thickness), maximum anatomical cross-sectional area (ACSA_max_), and fiber CSA. For this purpose, we recorded HD-sEMG signals, ultrasound, magnetic resonance imaging, and muscle biopsy of the biceps brachii muscle from two groups of participants – untrained-controls (UT=14) and strength-trained (>3 years of training, ST=16) – while they performed isometric ramp contractions with elbow flexors (at 15, 35, 50 and 70% maximum voluntary torque - MVT). We assessed the correlation between the number of accurately detected MUs by HD-sEMG decomposition and each measured parameter, for each target force level. Multiple regression analysis was then applied.

**Main results:** ST subjects showed lower MED (UT: 4.8 ± 1.4 vs. ST: 3.7 ± 0.8 mm) associated to a greater number of identified motor units (UT: 21.3 ± 10.2 vs. ST: 29.2 ± 11.8 MUs/subject). Both groups showed a negative correlation between MED and the number of identified MUs at low forces (r= −0.6, p=0.002 at 15% MVT). Moreover, the number of identified MUs was positively correlated to the distribution of MU territories (r=0.56, p=0.01) and ACSA_max_ (r=0.48, p=0.03) at 15% MVT. By accounting for all anatomical parameters, we were able to partly predict the number of decomposed MUs at low but not at high forces.

**Significance:** Our results confirmed the influence of subcutaneous tissue on the quality of HD-sEMG signals and demonstrated that MU spatial distribution and ACSA_max_ are also relevant parameters of influence for current decomposition algorithms.

## Introduction

Motor neurons discharge action potentials to muscles that translate these neural commands into forces and movement. A motor unit (MU) is a structure made of a motor neuron and its innervated muscle fibers. The summation of action potentials of the muscle fibers of a MU represents the motor unit action potential (MUAP). Therefore, the measured electromyographic (EMG) signal indirectly represents the output from the spinal cord [1]–[3]. The identification and tracking of individual MUs from EMG allows for the study of the neural code underlying movement [4]–[10].

Surface EMG (sEMG) is a non-invasive approach for acquiring neuromuscular activity by placing electrodes on the skin surface. The recent use of high-density grids of electrodes (HD-sEMG) provides sufficient spatial information to discriminate the action potentials of individual MUs [11]–[14]. The identification of active individual MUs is achieved through the combination of high-density grids for recording and decomposition algorithms for discriminating MUAPs [8], [15], [16].

Blind source separation (BSS) algorithms are widely used to decompose HD-sEMG signals. This approach identifies MU discharges with almost no prior knowledge. BSS-based EMG decomposition has been extensively validated on simulated signals and by comparison with the decomposition of intramuscular EMG recordings [17]–[20]. Although this approach has been proved reliable [21], there is a large variability in the number of identified MUs across subjects, with results ranging from 1 to ~30 detected MUs [7], [14]. Moreover, the number of identified MUs tends to decrease with increasing force levels [14].

The BSS procedure works well with HD-sEMG data because the sources, i.e. the series of discharge times of MUs, are sparse. However, decomposition accuracy also depends on the similarity of MUAP waveforms, which is influenced by the properties of the volume conductor. Although computer simulations can be used to investigate the influence of these factors on EMG decomposition [22], [23], an experimental analysis of the potential factors that can alter the number of identified MUs, including anatomical and neurophysiological differences across subjects, is still missing.

One factor of influence on MU decomposition is the degree of MU synchronization, which is associated to the correlation between sources [24]. Another factor of influence is related to the thickness and properties of the tissues between the muscle fibers and the recording electrodes (subcutaneous tissue). The subcutaneous layer acts as a low-pass filter, which reduces the differences in MUAP waveforms [25]–[28]. Since MUAP waveform shapes vary according to the MU location, the distribution of MU territories within the muscle could also impact the performance of decomposition algorithms. In addition, muscle size (i.e., anatomical cross-sectional area) and the cross-sectional area (CSA) of individual fibers could also affect the spatial distribution of the HDs-EMG data, due to the linear relation between individual muscle fiber diameters and amplitude of the MUAPs [29]–[32].

Therefore, in this study, we evaluated the impact of synchronization, distribution of MU territories, muscle size, fiber area, and thickness of the subcutaneous tissue on the decomposition of HD-sEMG data recorded from the biceps brachii (BB) muscle during voluntary isometric contractions. We specifically selected the BB muscle and two subject populations (sedentary and trained individuals) to maximize the ranges of variation of these factors.

We hypothesized that the analyzed factors had a strong influence on the ability of HD-sEMG algorithms to identify MU activity.

## Methods

### Participants

The data used in the present study was derived from a large dataset collection aimed at testing the neural control of movement in untrained healthy individuals versus a cohort of subjects that performed resistance training for more than 3 years. Results on this dataset have been recently published [33]. In the present study, we report data from HD-sEMG, magnetic resonance imaging (MRI), muscle biopsies, and ultrasound recordings for the total number of participants (30 healthy subjects). General requisites for participation in this experiment included: age between 18 and 40 years, no past upper limb surgery or traumatic injury, and no previous use of anabolic-androgenic steroids. The participants were assigned to two groups (UT - untrained and ST - strength-trained) based on specific criteria (N = 14 for UT and N = 16 for ST). The UT group did not perform any regular physical training or upper-body resistance exercise in the 18 months prior to the start of the study. Conversely, the ST group was involved in upper arm resistance training for at least 2 sessions a week for more than 10 months a year, for a minimum of 3 years. The ST group achieved a minimum of 90 Nm of maximum voluntary torque for isometric elbow flexion during the familiarization session.

The procedures were in agreement with the Declaration of Helsinki and approved by the Loughborough University Ethics Approvals Sub-Committee (R17-P174). All recruited subjects signed a written informed consent before they participated in this study.

### Overview of the study

The study was conducted in 3 sessions, in which the participants visited the laboratory. The first visit was introductory. The experimental procedures were explained, and the participants were familiarized with the experimental setup through the practice of the isometric sub-maximum and maximum voluntary contractions (MVCs) of the elbow flexors of the non-dominant arm that were to be performed in the main measurement session. They also answered questions about their physical activity habits by completing the International Physical Activity Questionnaire (IPAQ) [34]. Thus, based on the subjects’ strength and physical activity, their eligibility to this study was assessed.

The second and main measurement session (7-10 days after the first session) involved a first assessment of muscle-electrode distance (MED - subcutaneous fat layer thickness) with B-mode ultrasonography. Thereafter, following a standardized warm-up, this session consisted of the simultaneous recording of isometric elbow flexor force and HD-sEMG recordings from BB muscle during maximum voluntary contractions (MVCs) and submaximal ramp contractions at different force targets (15, 35, 50, 70% maximum voluntary torque - MVT). The third session occurred 2 or 3 days after the second one, it involved the collection of muscle biopsies – to assess the cross-sectional area of the muscle fibers; and the collection of magnetic resonance T1-weighted axial images along the length of the humerus to assess anatomical cross-sectional area (ACSA) of the BB. Participants were asked not to perform demanding physical exercise (48 h) and avoid caffeine consumption (24 h) before the main session.

### Experimental procedures

In the main session, the standardized warm-up consisted of 7 isometric contractions of the elbow flexors (3 x 50%, 3 x 75%, and 1 x 90% of self-perceived maximum voluntary force - 5s duration, 15-30s rest in between). Subsequently, the measurements of MVCs and ramp isometric contractions were performed, in the following order.

The participants performed 3 or 4 MVCs, of 3-5s duration, with 30 s of rest between the contractions. Verbal encouragement to achieve maximum force during each contraction was provided. The peak of force achieved was displayed on a monitor for real-time feedback to the participants. The highest force recorded during any of the MVCs was used as a reference to determine the intensity of the ramp isometric contractions.

After 5 minutes of rest, the participants were asked to match a ramp contraction template displayed on a monitor. They performed 8 ramp contractions (two at each force level) with a linear increase in force at 10% MVT/s from rest to the plateau of the contraction (either 15, 35, 50 and 70% MVT) that was then held steadily for 10s (for 15 and 35% MVT) or 5s (for 50 and 70% MVT). Each ramp contraction was separated by 3 to 5 min of recovery and performed randomly to minimize possible fatigue effects. The real-time force recording was superimposed to the template for visual feedback.

### Force recordings

For the first and second sessions, we used a custom isometric elbow flexion dynamometer. It comprises a rigid strength-testing chair that was adjusted according to participants’ height and upper arm length (both evaluated during the first visit). In the first session, a participant-specific configuration of the dynamometer was defined and reproduced for the subsequent session.

Participants were seated on the dynamometer with the hip joint flexed to 90°. Their waist, chest, and shoulder were firmly strapped to the back of the chair. The non-dominant shoulder joint angle was set at 90° of flexion with 10° of horizontal abduction with the posterior of the upper arm resting on a horizontal board. The elbow joint angle was set at 70° of flexion, and the forearm was in half-supination 45° [35], [36]. The non-dominant wrist was strapped to an adjustable wrist brace that was in series with an S-beam tension-compression strain gauge (Force Logic, Swallowfield, UK) fixed perpendicular to the forearm. The contralateral arm rested on the thigh. The force signal from the strain gauge was amplified (x 200) and sampled at 2,048 Hz by an external analog to digital converter - 16-bit A/D (EMG-Quattrocento, OT Bioelettronica, Turin, Italy). The force signal was recorded with the OTBioLab software (OT Bioelettronica). Force and ramp contraction template feedback were provided by the Spike 2 software (CED, Cambridge, UK).

### HD-sEMG signal recordings

In order to position the electrodes correctly, the muscle bellies (BB long and short head) were identified by an experienced investigator, and the ultrasound was used to mark the profiles of the muscle. The BB perimeter was marked with a surgical marking pen. In this area, the skin was shaved, lightly abraded, and cleansed with 70% ethyl alcohol. After the initial configuration of the dynamometer, 2 grids of 64 electrodes each (13 rows x 5 columns - 10.9 cm x 3.7cm; gold-coated; 1-mm diameter; 8-mm interelectrode distance - IED; OT Bioelettronica) were positioned over the surface of the BB muscle (non-dominant arm). The two grids were aligned to form an array of 128 electrodes covering the long and short heads of the muscle (Figure 1), with the electrode arrays placed with their center on the muscle bellies and aligned with the direction of the muscle fibers.

**Figure 1.**
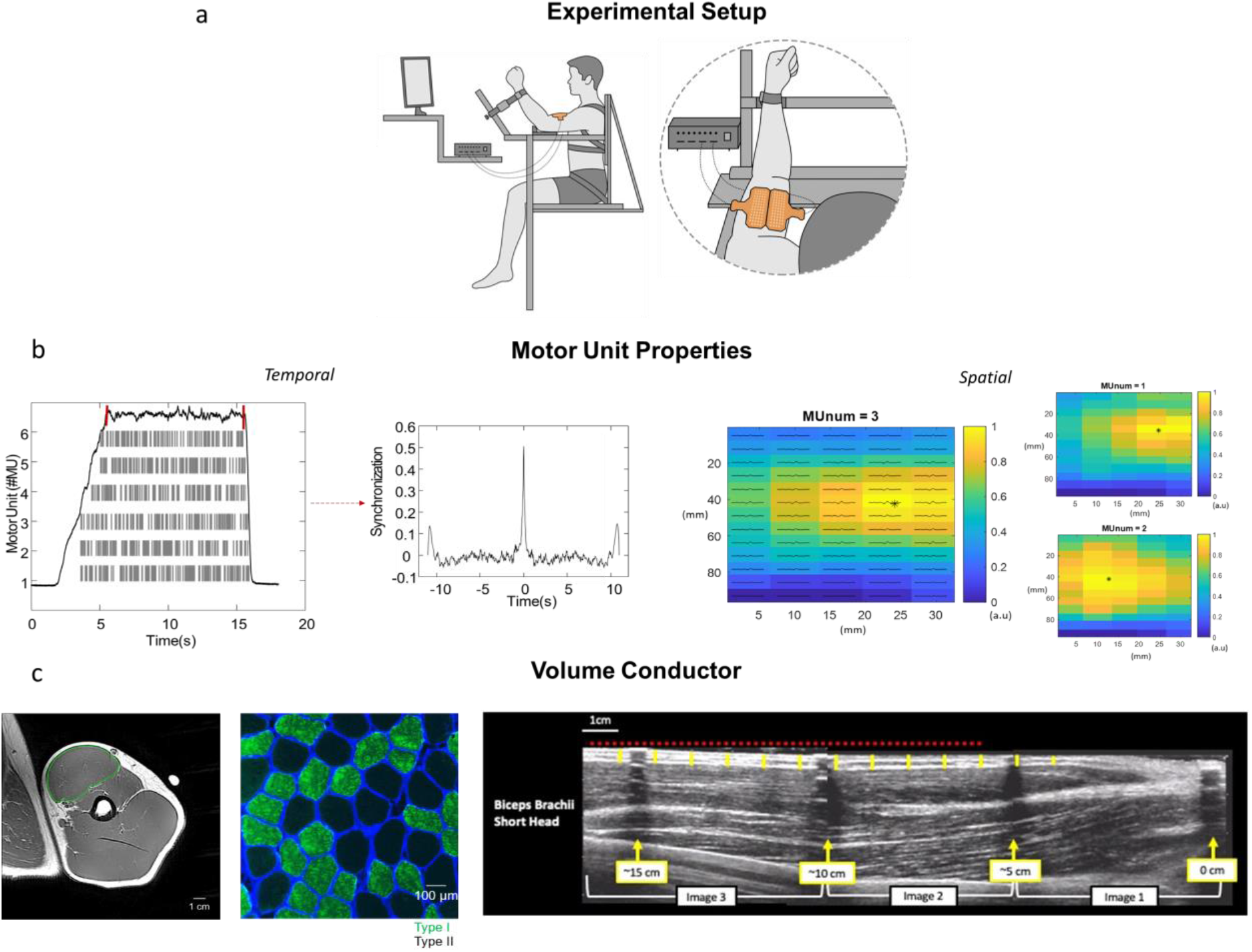
Overview of the study (a) Experimental setup and electrodes’ location on the biceps brachii (BB) muscle (b) motor unit (MU) properties extracted from the study, from left to right: motor unit synchronization and spatial distribution (c) Volume conductor parameters extracted from the study, from left to right: maximum anatomical cross-section (ACSA_max_), cross-sectional area of individual fibers (fiber CSA) and muscle-electrode distance (MED - proportional to subcutaneous fat thickness).

To improve the electrode-skin contact, a disposable bi-adhesive foam layer was aligned to the electrodes, its holes were filled with conductive paste (SpesMedica, Battipaglia, Italy) and attached to the skin. The reference electrodes for each electrode grid were positioned over the radial styloid process. The main ground electrode was placed on the styloid process of the ulna.

The analog HD-sEMG signals were recorded in monopolar mode (amplified 150x), sampled at 2,048 Hz, bandpass filtered 10-500Hz, and converted to digital signals using a multichannel amplifier 16-bit A/D (EMG-Quattrocento). HD-sEMG signals were recorded with the software OTBioLab and synchronized at the source with the force signal by the same acquisition system. The offline analysis was performed in MATLAB (MATLAB R2021a, Mathworks, Natick, MA).

### Ultrasound measurements

The subcutaneous fat thickness is obtained as an indirect measure of the muscle-electrode distance (MED) that consists of subcutaneous fat layer, skin, and superficial aponeuroses of the muscle.

The MED along the length of the BB long head and short head from the non-dominant arm was measured using a B-mode ultrasonography machine (EUB-8500, Hitachi Medical Systems UK Ltd, Northamptonshire, UK). During ultrasound image capture participants stood upright with their shoulder joint abducted to ~90° (relative to the anatomical position, i.e., humerus horizontal) with the elbow extended and the palm of their hand supinated. The upper arm and elbow were supported by adjustable padding on top of the monitor of the ultrasound machine and participants were instructed to stand still and relax their arm and shoulder so that the arm musculature was at rest during image capture. The borders of the BB long and short head were marked and a center line running along the length of each muscle at 50% of the medio-lateral width was marked with indelible ink. Narrow Echo-absorbant markers (layers of transpore tape; Figure 3b) were placed on the dermal surface perpendicular to the length of the humerus on the distal muscle-tendon junction of the BB (i.e., 0 cm) and at 5 cm intervals (5, 10, 15 cm) along the length of the BB (i.e., covering both short and long heads with each echo absorbant marker).

**Figure 2.**
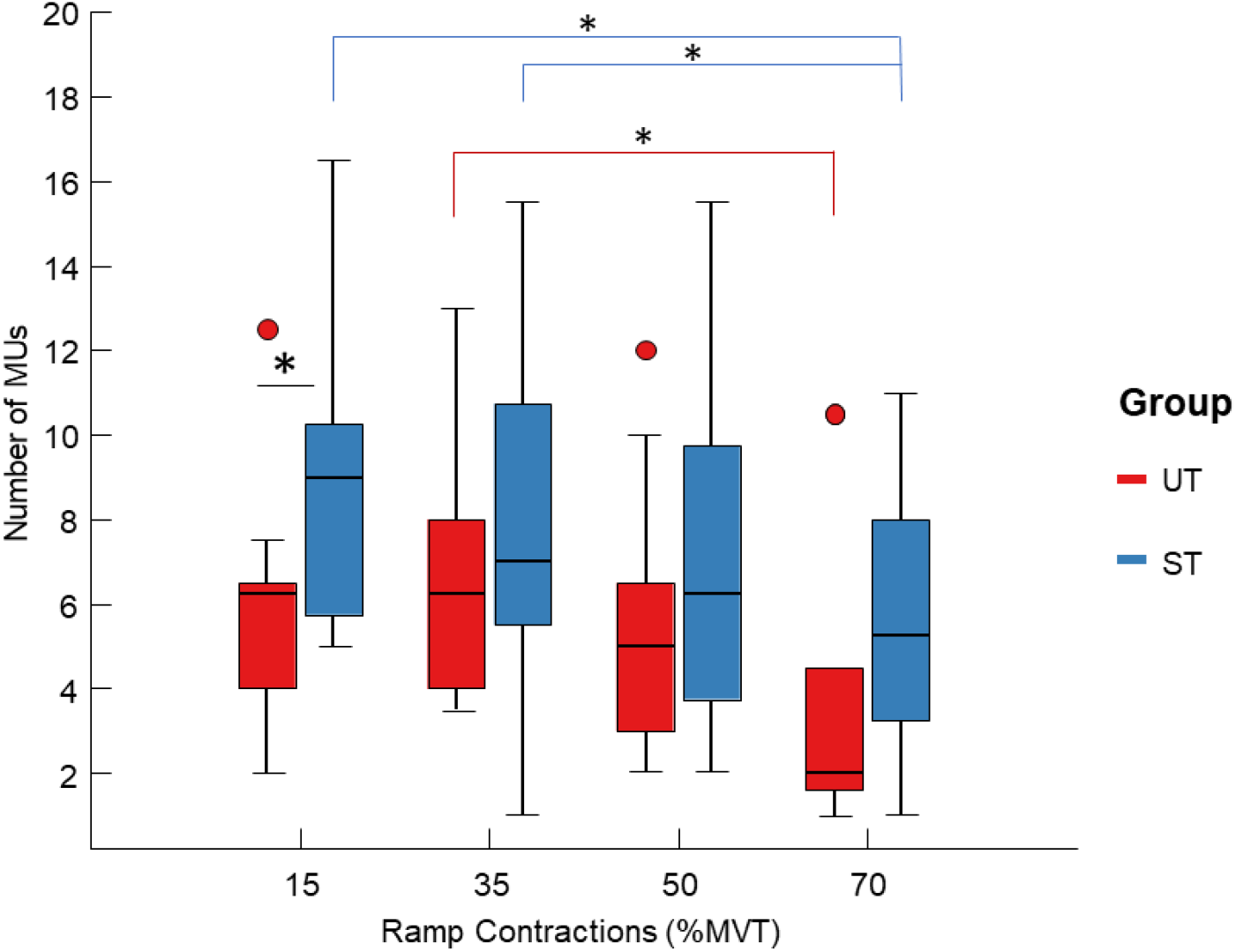
Boxplot of the number of identified motor units (MUs) for both subject groups: Untrained - UT (in red) and Strength Trained - ST (in blue), according to target forces (15, 35, 50, and 70% maximum voluntary torque - MVT). Red dots indicate outliers. * 0.01 < *p-value* < 0.05; **0.001 < *p-value* < 0.01.

**Figure 3.**
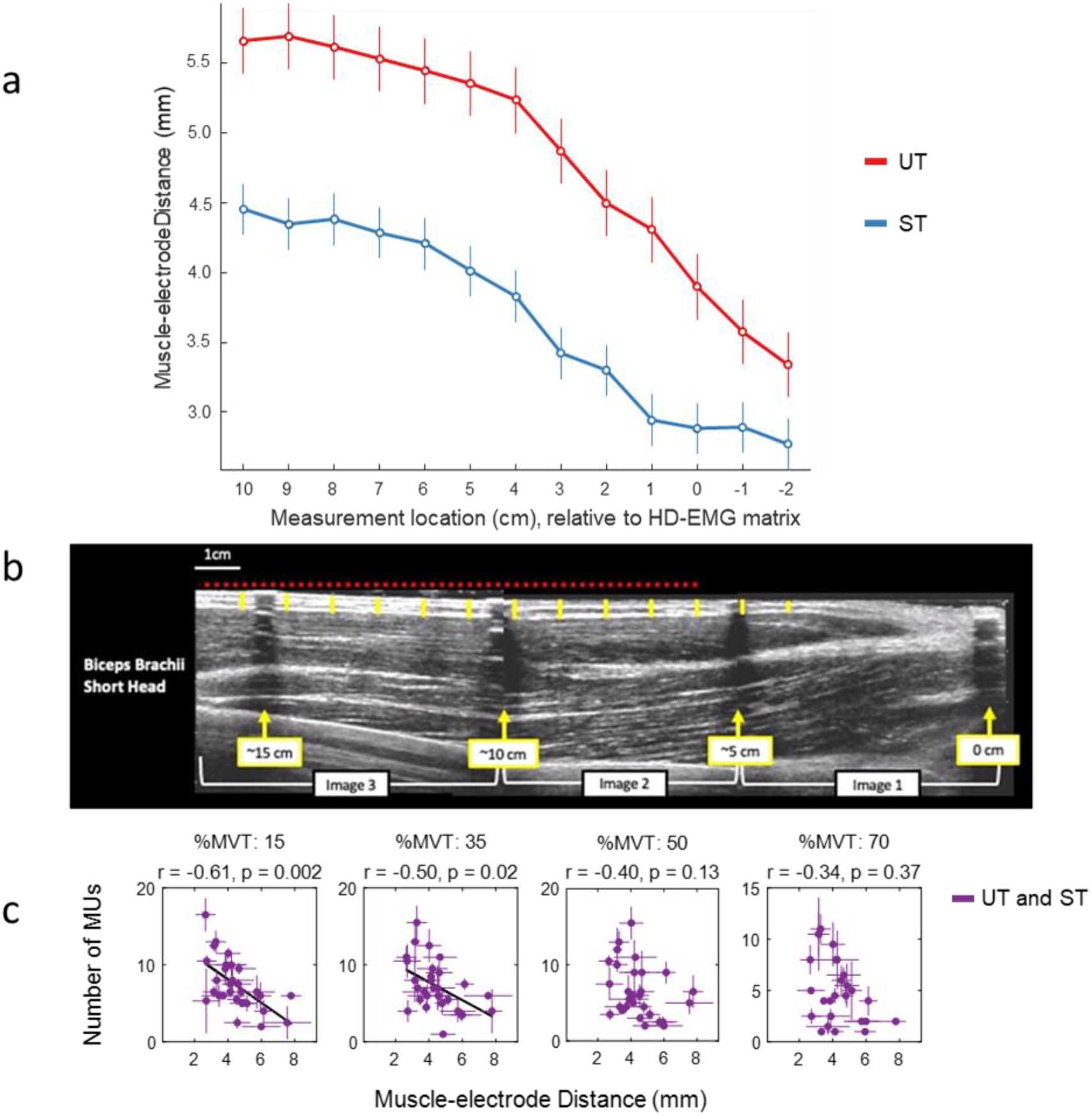
(a) Average of muscle-electrode distance (MED)/subcutaneous thickness across all participants, according to their measurement location. (b) Ultrasonography images of the biceps brachii (BB) short head, captured as a digital video recording with an investigator gradually moving the ultrasound probe along the muscle on a predetermined line marked on the dermal surface. The MED (yellow vertical lines) was measured at 13 continuous 1 cm intervals along the length of the muscle relative to the position of the high-density EMG array (denoted as the red-dashed horizontal line, 0cm position is the distal edge of the array). (c) Correlation between the number of identified motor units (MUs) and the averaged muscle-electrode distance, according to force levels (15, 35, 50 and 70% maximum voluntary torque - MVT), for both groups combined, Untrained - UT and Strength Trained - ST (in purple).

Water soluble transmission gel was applied along the center line of each BB muscle to ensure ultrasound image detection was possible along the entire length of each muscle. Then a 92-mm 5-10 MHz linear-array transducer (EUP-L53L; also coated with transmission gel) was gradually (over ~30 seconds), and moved with minimal pressure along the center line of each muscle from the distal BB tendon to the proximal end of each muscle. A single sweep of each muscle was performed, unless image quality was deemed unacceptable in which case a second sweep was performed. Video output from the ultrasound machine was transferred to a computer (via an S-video to USB converter), and the ezcap video capture software was used to record images. Images were subsequently imported into public domain software and analyzed (Tracker, version 5.0.6 http://www.physlets.org/tracker/).

The Echo-absorbant markers were used to align three separate images (0-5 cm, 5-10 cm, and 10cm+) and allow MED to be measured at continuous 1 cm intervals along the length of each BB muscle. Specifically, MED was measured as the distance from the surface of the skin to the muscle fascia at thirteen 1-cm intervals along each muscle (Figures 1 and 3b), starting −2-cm distal relative to the placement of the distal row of electrodes of the HD-sEMG grid, and then every 1cm until 13 cm above the start location given array length of 10.9 cm. This approach was taken to at least partially account for distal movement of the underlying muscle beneath the HD-sEMG grid between the ultrasound measurement (i.e., elbow full extended) and elbow flexion contractions (i.e., elbow flexed to 70°).

### Muscle size

T1-weighted axial images of the non-dominant arm were obtained using a 3T MRI scanner (Discovery MR750w, GE Healthcare, Chicago, IL, USA) and a receiver 16-channel flex coil. All the scans were performed with the subjects in a supine position, the elbow joint fully extended and relaxed. The axial images were taken in three overlapping blocks, from the humeral head to below the elbow joint – aligned with the humerus. The scanning parameters were the following: repetition time = 600ms, time to echo = 12.8ms, field of view = 180×180mm, image matrix = 260×260, pixel size = 0.69×0.69mm, slice thickness = 5mm and interslice gap = 5mm - PROPELLER mode.

The anatomical cross-sectional area (ACSA) of the BB muscle was segmented (as one mass for long and short heads) along the humerus using a public domain DICOM software (Horos, version 3.3.6, www.thehorosproject.org). The measurement of maximum ACSA (ACSA_max_) of the BB muscle was used as a metric of muscle size.

### Muscle biopsy and fiber analysis

Muscle biopsies were taken from the biceps brachii of the non-dominant arm under local anesthesia (1% lidocaine) using a conchotome biopsy needle as previously described [37]. Muscle samples were dissected of any visible fat and connective tissue and blotted dry and then immediately embedded in a mounting medium (Tissue-Tek O.C.T. Compound, Sakura Finetek Europe, Alphen aan den Rijn, The Netherlands) and frozen in liquid nitrogen-cooled isopentane and stored at −80°C for immunohistochemistry analysis.

Transverse serial sections (8 μm) were obtained using a cryotome and placed onto poly-L-lysine-coated glass slides. Sections were fixed for 10 min in 3.7% paraformaldehyde at room temperature and blocked with tris-buffered saline containing 2% bovine serum albumin, 5% goat serum, and 0.2% Triton for 1 h at room temperature. Serial muscle sections were then incubated with a primary antibody for myosin heavy chain I (A4.951, Developmental Studies Hybridoma Bank, Iowa City, USA) diluted 1:200 in the blocking solution for 1 h at room temperature. Sections were then incubated for 2 h at room temperature with an appropriate secondary antibody consisting of goat anti-mouse Alexa Fluor 488 (A11029, Fisher Scientific, Pittsburgh, USA) diluted 1:500 and wheat germ agglutinin Alexa Fluor™ 350 Conjugate (W11263, Fisher Scientific, Pittsburgh, USA) diluted 1:20 in the blocking solution. Following incubation, coverslips were mounted with Fluoromount aqueous mounting medium (F-4680, Sigma-Aldrich, St. Louis, USA).

Images were captured using a fluorescence microscope (Leica DM2500) at x20 magnification (Figure 1). Image analysis was undertaken using Fiji (ImageJ) software. The investigator was blinded to the participant code of each sample. Only transversely sectioned fibers were included in the analysis, i.e., any fibers that were clearly oblique or not transverse to the long axis of the fiber were excluded. Fiber cross-sectional area was assessed by manually drawing around the perimeter of each fiber for 200 different fibers per participant. In 4 participants (3 in ST and 1 in UT), only 145-165 fibers were analyzed for area, given the insufficient number of clear images/fiber perimeters. In addition to the fibers analyzed for area, a total of 500 fibers per participant were counted as type I or type II fibers. Fiber type composition was expressed in two ways: 1) percentage by fiber number i.e., the number of fibers of each type relative to the total number of fibers counted (n = 500) and 2) percentage by fiber area i.e., the summed area of the fibers of each type relative to the total area of fibers analyzed (n = 200). We used the total sum of fibers cross-sectional area (fiber CSA) for our analysis.

### Data Analysis

#### Force

The force signals were analyzed offline in MATLAB. First, the signal was converted to Newtons (N), and the offset was removed by subtraction of the force baseline. A low-pass Butterworth filter (fourth-order, zero lag, 15-Hz cut-off frequency) was applied to the signal. The torque (Nm) was calculated by multiplying the force (N) by the lever arm length (distance from the elbow joint center to the center of the wrist brace). The isometric ramp contractions at all force levels and all trials were analyzed (15, 35, 50, and 70% MVT).

The absolute value (N) and normalized force values were determined by averaging the force values throughout the entire plateau phase of the isometric ramp contractions at the target force levels.

#### HD-sEMG

The recorded HD-sEMG monopolar signals were band-pass filtered (Butterworth filter, second-order, zero lag, 20 and 500-Hz cut-off frequencies). The filtered signals were decomposed into individual MUs through a convolutive BSS algorithm [19], [38]. The decomposition accuracy was assessed by calculating the pulse-to-noise ratio (PNR) for each MU [38]. The discharge times of the MUs were identified and manually inspected by an experienced investigator [14]. Only the MUs with PNR > 30dB and interspike interval < 2s were kept [17], [38] for the visual inspection. Only data from the medial grid (BB shorthead) was processed and analyzed because it presented a greater number of MUs in comparison to the lateral grid (BB long head).

#### Motor unit synchronization

From the discharge times of the identified MUs obtained by decomposition, we extracted the spike trains that occurred during the force plateau of the ramp contractions. Thereafter, a moving-average filter (Hanning window of 100 ms duration) was applied to the spike trains [39]. The MUs were randomly arranged in pairs to obtain a normalized correlation value that was not dependent on the number of identified MUs. The cross-correlation of each pair was computed, and the results were averaged across 100 random permutations [39], [40]. The maximum value of this average was defined as a synchronization metric.

#### Motor unit spatial distribution

MUAPs were extracted through spike-triggered averaging of HD-sEMG signals. Based on the RMS values from the MUAPs, we generated a spatial map for each MU. In order to estimate the MU location, we computed the weighted centroid of each MUAP according to their high activity region (it is possible to have more than one high activity region per MU). For this, we used the image processing toolbox of MATLAB. First, as an intermediate step, the MUAP maps were converted into grayscale from 0 to 1 (0-black, 1-white). For the weighted centroid calculation, a region with RMS values higher than 0.8 was selected, from which a weighted average of the pixels coordinates was performed, with the pixel intensities (grayscale) as weights.

Using the weighted centroid coordinates of each MUAP (*x_i_*, *y_i_*) with *i* = *1*, *2*,…, *n*, being the number of MUs, we established a metric to evaluate the spatial distribution of the MU centroids for each participant and force level (e.g., subject 1, 35% MVT). The metric chosen was the standard distance, 2D analogous of the standard deviation [Equation (1)]. First, we calculated the mean center of the weighted centroids (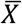, *Ȳ*) and the Euclidean distance between the weighted centroids and the mean center (see Figure 4) for each MU. The standard distance (*Stdist*) was defined as follows:

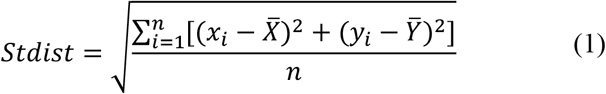

**Figure 4.**
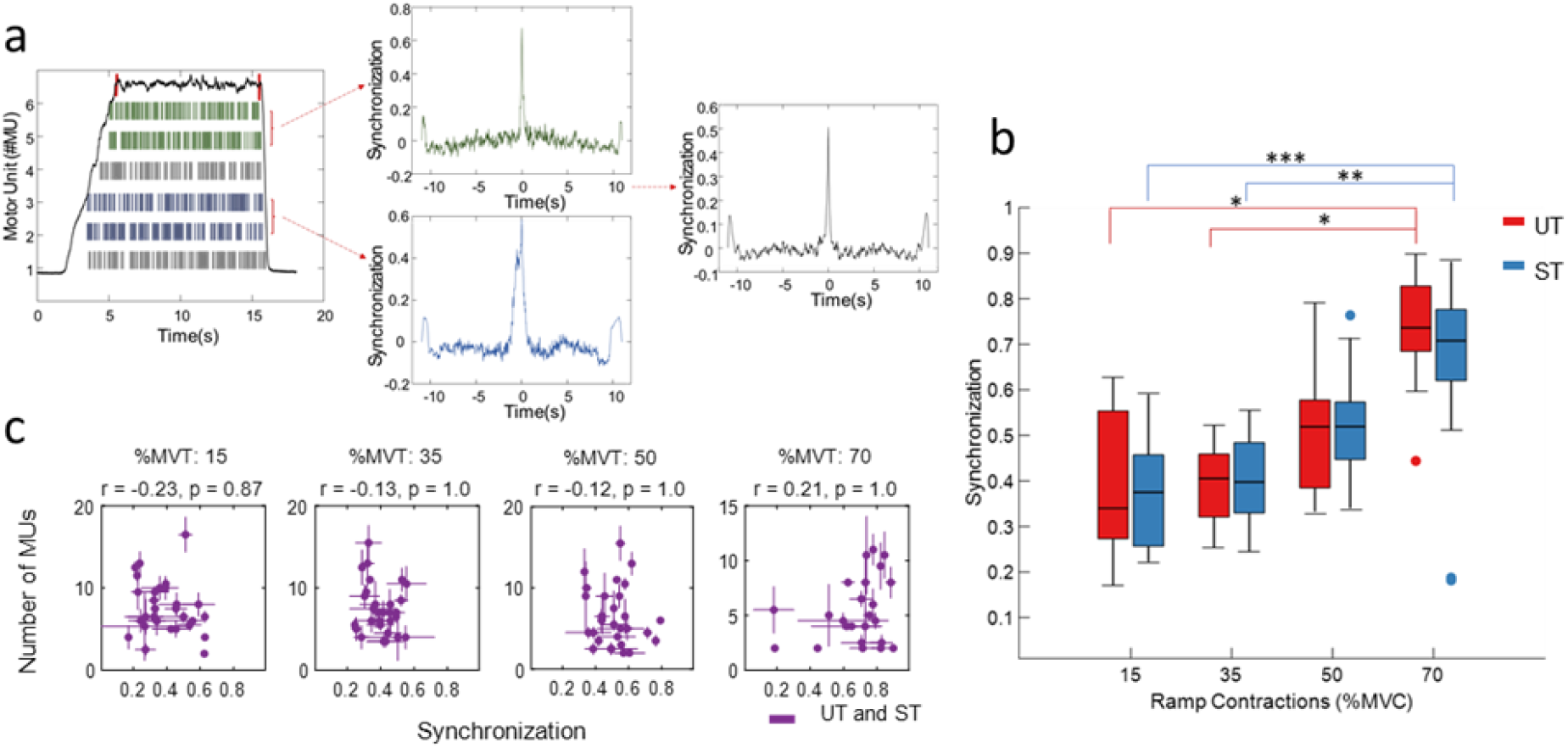
(a) Calculation of motor units’(MUs) synchronization. MUs were arranged in random pairs, the synchronization for each pair was calculated and averaged after 100 random permutations. The maximum value of this average was used. (b) Boxplot of MUs synchronization for both groups Untrained (UT - in red) and Strength Trained (ST - in blue), according to target forces (15, 35, 50 and 70% maximum voluntary torque - MVT). Red and blue dots indicate outliers. * 0.01 < *p-value* < 0.05; **0.001 < *p-value* < 0.01; *** *p-value* < 0.001. (c) Correlation between the number of identified MUs and synchronization, according to contraction levels, for both groups (in purple).

The standard distance is, therefore, a measure normalized by the number of MUs (*n*). It expresses the spread of the centers of the MU territories in the horizontal skin plan.

### Overall Analysis and Statistics

Since there were two trials for each subject for the same force target (e.g., 2 x 50% MVT), we first averaged the number of MUs, MUs synchronization, and standard distance (distribution of MU territories) between the trials. The data were organized according to the force targets (15, 35, 50, and 70% MVT). We investigated the influence of the target forces on the number of identified MUs, MUs synchronization, and standard distance, separately. Before the analysis, we assessed the normality of the data using a Shapiro Wilk test. Because most of the variables did not present a normal distribution, we adopted a non-parametric approach for statistical comparisons.

To evaluate whether there were differences between groups (UT vs. ST) of same % MVT, a Mann-Whitney U test was applied (significance p-value < 0.05). A Friedman test was applied (analogous to the parametric repeated measures ANOVA) to assess the difference within the groups, followed by pairwise comparisons. Bonferroni correction was performed when the results were significant.

Spearman correlation (p-value < 0.05, Bonferroni correction) was used to determine the relationship between the number of MUs with MU synchronization and standard distance. In this case, we considered only the recordings with two or more MUs. Regarding the MED, we averaged it across subjects according to the same measurement location along the arm length/electrode grid. For each subject, we averaged the MED from the measurement points 0 to 10 cm (according to the HD-sEMG grid position – 0 cm being the distal edge of the grid). We opted for measures that were inside the area delimited by the HD-sEMG grid. The ACSA_max_ and sum of fiber CSA were also correlated with the number of MUs. Lastly, a multiple regression analysis was used to investigate the relationship between the dependent variable - number of MUs - and the predictor variables - MED, synchronization, standard distance, ACSA_max_, and sum of fiber CSA. A box-cox transformation was applied when the data were not normally distributed, or in general, when they did not satisfy multiple regression assumptions (e.g., heteroscedasticity, kurtosis). The individual contribution of each of the parameters to the variance in the number of MUs was assessed [41], [42]. Multiple regression results are reported including F-statistic and degrees of freedom, β - standardized coefficient, t, and p-values. A backward stepwise regression was also performed to verify if any parameter could be removed from the model, improving it significantly.

The statistical analyses were performed with MATLAB and the software RStudio (RStudio: Integrated Development Environment for R, Version 2021.9.1.372, RStudio, PBC, Boston, MA).

## Results

With the aim to understand the variability in the number of decomposed MUs from HD-sEMG signals found in different subjects within the same tested muscle, we extracted some potential parameters that might affect the spatiotemporal representation of the MUAPs: MED, MU synchronization, MU spatial distribution, maximum anatomical cross-sectional area (ACSA_max_) and sum of fiber cross-sectional area (fiber CSA) from the BB muscle. We then evaluated their influence on the number of identified MUs via correlation analyses. An overview of the study is presented in Figure 1.

Considering all the 30 participants, averaging the trials across the same subjects at any level of contraction (15, 35, 50, 70% MVT), resulted in UT = 298 and ST = 468 MUs, with an average of 21.3 ± 10.2 MUs/subject for the UT group, and of 29.2 ± 11.8 MUs/subject for the ST group. The average number of MUs for each target force, considering all the trials, is shown in Table 1.

**Table 1.**
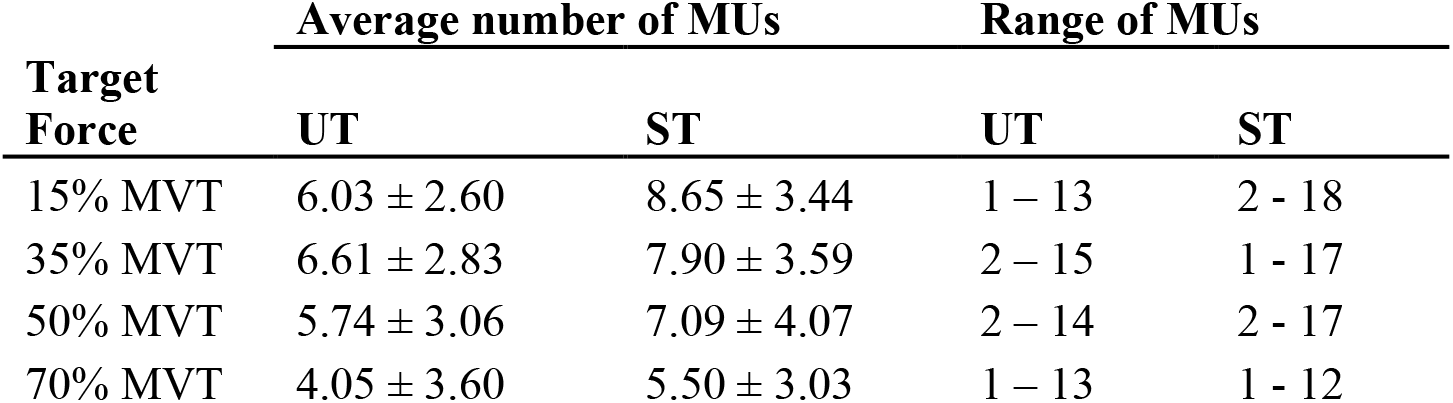
Description of the variability in the number of identified MUs across target forces (15, 35, 50 and 70% MVT - maximum voluntary torque), considering all trials. The average number of MUs is shown as mean ± SD. UT – Untrained group, ST – Strength trained group.

| Target Force | Average number of MUs |  | Range of MUs |  |
| --- | --- | --- | --- | --- |
|  | UT | ST | UT | ST |
| 15% MVT | 6.03 ± 2.60 | 8.65 ± 3.44 | 1 – 13 | 2 – 18 |
| 35% MVT | 6.61 ± 2.83 | 7.90 ± 3.59 | 2 – 15 | 1 – 17 |
| 50% MVT | 5.74 ± 3.06 | 7.09 ± 4.07 | 2 – 14 | 2 – 17 |
| 70% MVT | 4.05 ± 3.60 | 5.50 ± 3.03 | 1 – 13 | 1 – 12 |

The distribution of the average number of identified MUs for both groups (UT and ST) according to force level (15, 35, 50 and 70% MVT) is shown in Figure 2. In both Table 1 and Figure 2, we observed a higher number of MUs amongst the ST group. For 70% MVT, particularly for the UT group, a lower number of MUs was found, which is justified by an increased number of superimposed MUAPs in the EMG signal at higher force levels.

A significant between-group difference was only observed at 15% MVT (*p* = 0.02 – Mann Whitney U test). Within groups, there was a significant difference between MU number distributions at 70% MVT and 35% MVT (Friedman test: *p* = 0.005, *p*_35-70%_ = 0.006). For the ST group, the 70% MVT MU distribution differed significantly from 15 and 35% MVT distributions (Friedman test: *p* = 7 x10^-4^; *p*_15-70%_ = 0.006, *p*_35-70%_ = 0.002). This statistical difference between 70% MVT and other force levels is the result of its lower number of MUs.

### Muscle-electrode distance

We averaged the MED (proportional to the subcutaneous fat thickness) across all subjects along the muscle length relative to the distal edge of the HD-sEMG array (Figure 3a). The HD-sEMG grid was positioned according to the BB bellies profiles outlined using the ultrasonography. The distal edge of the array corresponds to the 0-cm position, and muscle-electrode measures were taken at thirteen 1-cm intervals along the muscle (Figure 3b), starting at −2-cm relative to the distal row of electrodes of the HD-sEMG grid.

The UT group presented a higher subcutaneous thickness compared to the ST group (mean ± SD: UT = 4.8 ± 1.4 mm; ST = 3.7 ± 0.8 mm), as shown by its higher MED (Figure 3a). For each subject, we averaged the MEDs from 0 to 10 cm along the BB to be correlated with the number of identified MUs.

Additionally, we assessed the correlation between the averaged MED(x-axis) and the number of identified MUs (y-axis) for the combination of UT and ST groups (Figure 3c). We noted a significant negative correlation with the MED at the force levels of 15% and 35% MVT.

As expected, our findings show that the subcutaneous fat acts as a low pass filter for the EMG signals. Regarding the ST group, a reduced subcutaneous thickness (Figure 3a) could explain why, in general, an increased number of identifiable MUs was found for this group (Figure 2).

### Motor unit synchronization, fiber cross-sectional area and pulse to noise ratio

We observed a higher synchronization of MUs for 70% MVT in both groups (Figure 4b). MU synchronization increased with voluntary force, although at 15% MVT (UT) it presented a larger range of values. No significant difference was found between the group’s synchronization distributions. Within the same group, the synchronization of MUs at 15 and 35% MVT was significantly different than for 70% MVT (Friedman test: UT – *p_15-70%_* = 0.04, *p_35-70%_* = 0.01; ST – *p_15-70%_* = 0.0003, *p_35-70%_* = 0.002), confirming an increased synchronization at this force level.

In order to assess whether the number of identified MUs across subjects is dependent on the synchronization level, a correlation analysis was performed (Figure 3c). No significant linear correlation was found between groups or for any force target. This result indicates that MUs synchronization may not affect the number of identified MUs across individuals.

Analogous to synchronization, we did not find any associations between muscle fiber CSA (obtained from the muscle biopsies) and the number of detected MUs, implying that MUAP amplitudes are not directly affecting MU decomposition. Also, the pulse-to-noise ratio was not correlated with any of the variables extracted in the study.

### Motor unit spatial distribution

We estimated MU centroids across the HD-sEMG grids to verify the distribution of MU territories in this skin plane and how it would affect the identification of MUs. We aimed at inspecting the differences between MUAPs, evaluating whether the MUAPs have more distant centroids when more MUs are detected.

Figure 5 shows an overview of the estimated distance across identified MUs. The standard distance across force levels (i.e., the distribution of MUs in space, Figure 5b), decreased with an increase in voluntary force. However, this was not observed for the ST group. No significant between-group difference was found. Within groups, for the UT group, the 70% MVT standard distance was significantly different than for 15 and 35% MVT (Friedman test: *p* < 0.001; UT – *p_15-70%_* = 0.02, *p_35-70%_* = 0.0004). For the ST group, the 50% MVT standard distance differed significantly than for 15 and 35% MVT (Friedman test: *p* = 0.002; ST – *p_15-50%_* = 0.02, *p_35-50%_* = 0.008).

**Figure 5.**
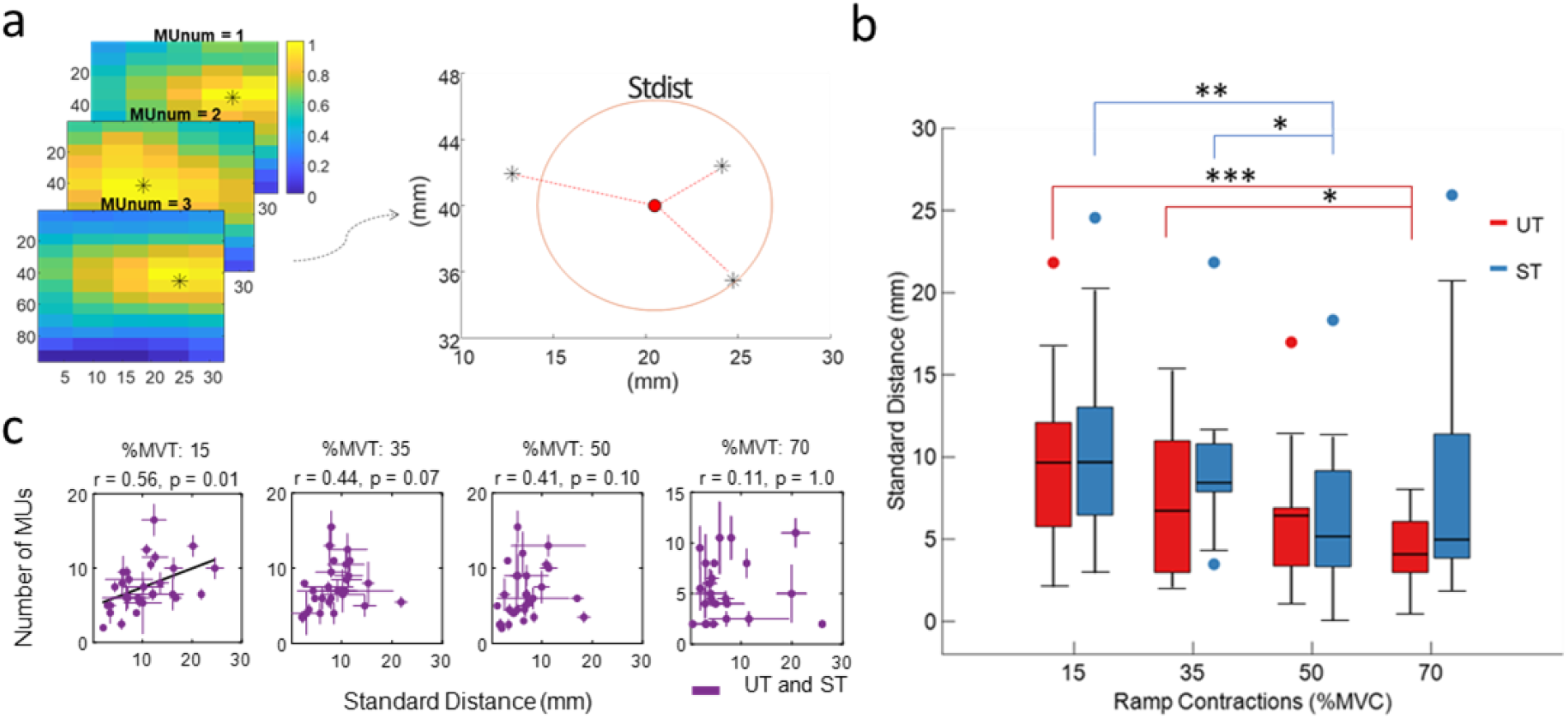
(a) Calculation of standard distance (stdist). The weighted centroid coordinates (indicated by *) of each motor unit action potential (MUAP) were obtained, and for each subject and force level the standard distance was calculated, with respect to the mean center of the MUAPs. (b) Boxplot of standard distance for both groups, Untrained (UT - in red) and Strength Trained (ST - in blue), according to target forces (15, 35, 50 and 70% maximum voluntary torque - MVT). Red and blue dots indicate outliers. * 0.01 < *p-value* < 0.05; **0.001 < *p-value* < 0.01; \*\*\**p-value* < 0.001. (c) Correlation between the number of identified MUs and the standard distance, according to contraction levels, for both groups combined (in purple).

We also evaluated the correlation between the standard distance and the number of identified MUs, considering both UT and ST groups together (Figure 5c). We noted a moderate positive linear correlation for 15, 35 and 70% MVT. This correlation was significant only at 15% MVT and marginally significant at 35 and 70% MVT.

### Anatomical cross-sectional area

The ST group showed a higher ACSA_max_ when compared to the UT group (ST = 18.4 ± 2.7 cm^2^; UT = 10.7 ± 2.1 cm2). The correlation between ACSA_max_ of BB and the number of detected MUs was tested (Figure 6). A significant positive correlation was found only at 15% MVT. For the other force levels, we also observed a positive correlation, although non-significant. Overall, our results indicate that the number of identified MUs increased with muscle size.

**Figure 6.**
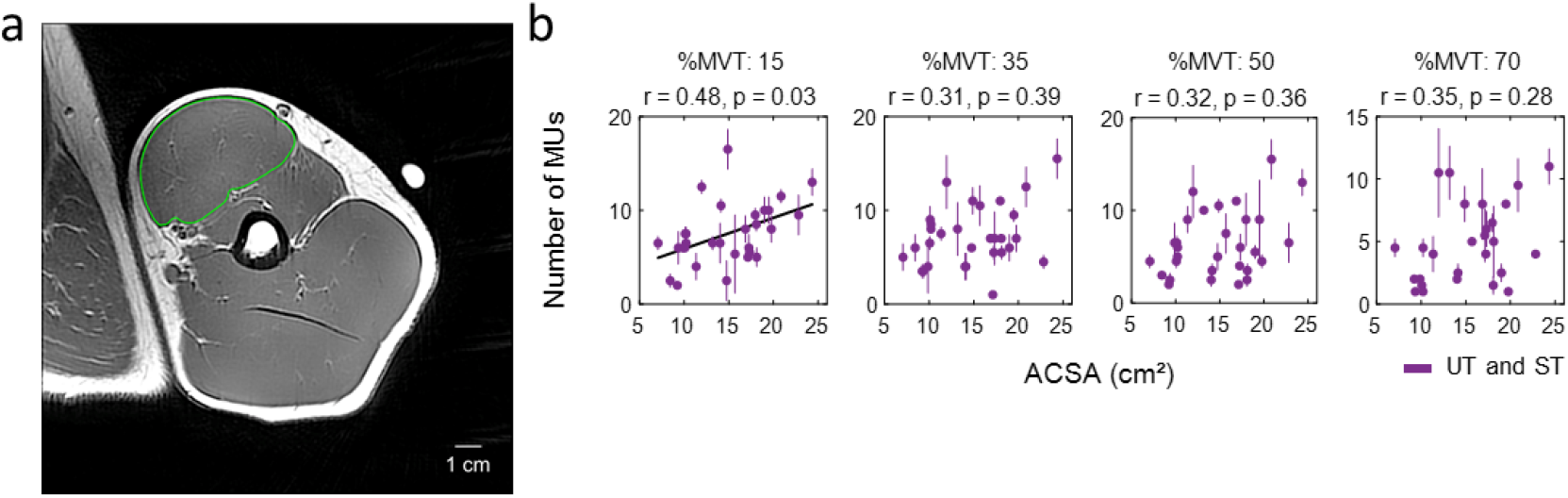
Correlation between the number of identified motor units (MUs) and the maximal anatomical cross-sectional area (ACSA_max_) of BB muscle, according to contraction levels (15, 35, 50, and 70% maximum voluntary torque - MVT), for both groups combined, Untrained - UT and Strength Trained - ST (in purple).

### Multiple regression model

A multiple regression was performed to model the relationship between the number of detected MUs (dependent variable) and the previously analyzed parameters (MU synchronization, MU spatial distribution, MED, ACSA_max_, and fiber CSA). We obtained two multiple regression models, according to the target forces that were grouped into low (15 and 35% MVT) and high forces (50 and 70% MVT).

At low forces, the model explained 44.3% of variance in the number of MUs (R^2^ = 44.3%; Adjusted R^2^ = 38.6%) and the regression was statistically significant (F (5,49) = 7.8, *p* < 0 .001). At higher forces, the model explained 26.3% of variance in the number of MUs (R^2^ = 26.3%; Adjusted R^2^ = 18.1%) and was also significant (F (5,45) = 3.2, *p* = 0.015).

MED was a significant predictor at low forces (low forces: *β* = −0.37, *t* = −3.00, *p* = 0.004; high forces: *β* = −0.19, *t* = −1.470, *p* = 0.15), followed by the MU spatial distribution (Low forces: *β* = 0.27, *t* = 2.45, *p* = 0.02; higher forces: *β* = 0.22, *t* = 1.84, *p* = 0.07). Figure 7 presents the relative contribution of each variable to the number of identified MUs, considering the R^2^ of the multiple regression. The contributions of MED and MU spatial distribution were, respectively, 19.3% and 13.4%. While at high forces, this percentage was reduced to 8.3% and 7.9%, with a marginal significance. ACSA_max_ (low forces: *β* = 0.18, *t* = 1.43, *p* = 0.16; high forces: *β* = 0.24, *t* = 1.72, *p* = 0.09) represented a contribution of 8.6% and 8.8% at low and high forces, respectively. Synchronization and sum of fiber CSA had the lowest contributions and could be eliminated from the model after a backward stepwise regression.

**Figure 7.**
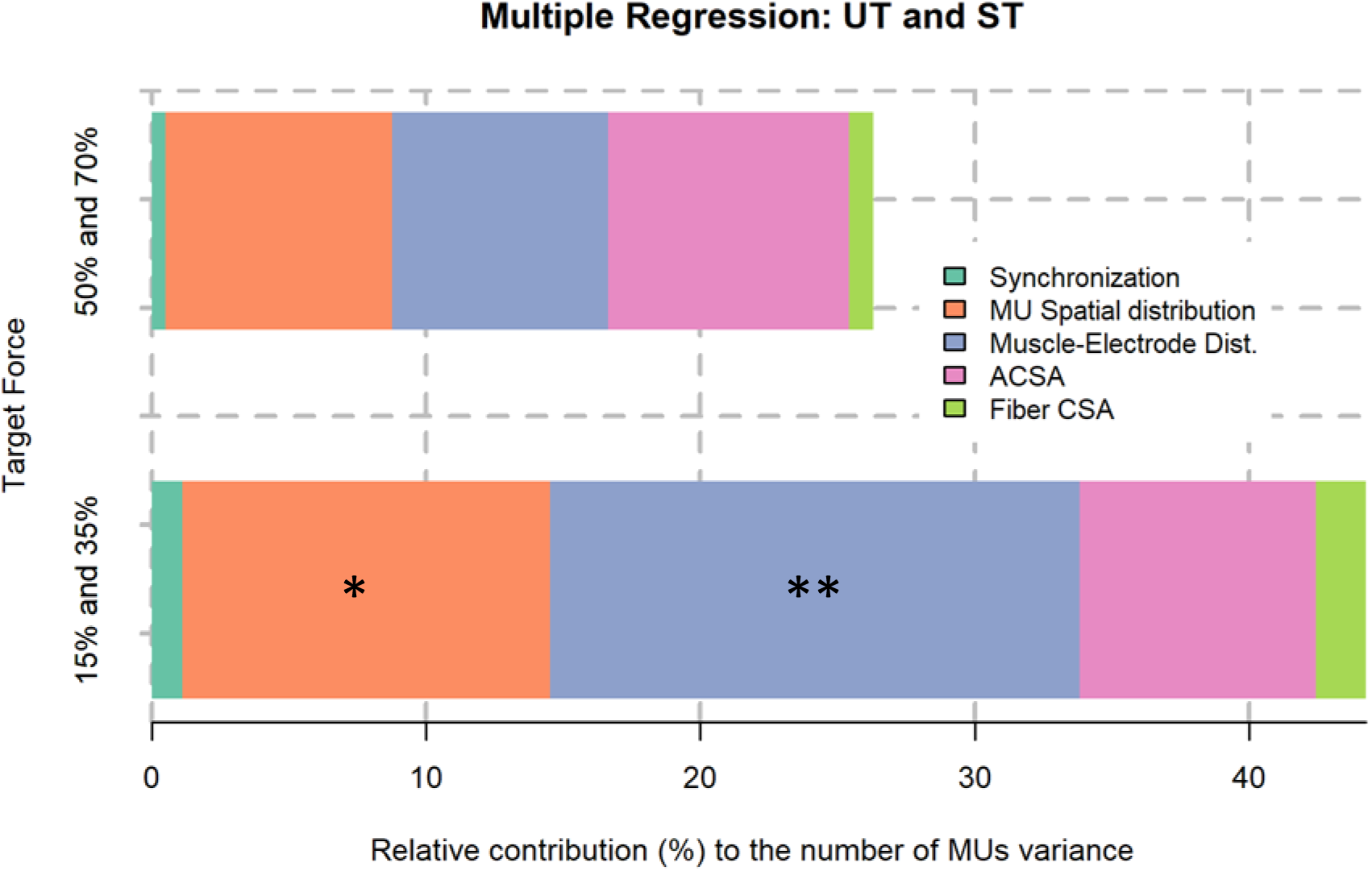
Relative contribution of predictor variables to R^2^ of multiple regression model, explaining the variance in the number of identified motor units (MU s). R^2^ (15 and 35% - maximum voluntary torque - MVT) = 44.3%; R^2^ (50% and 70%) = 26.3%. * *0.01* < *p-value* < *0.05*; \*\**0.001* < *p-value* < *0.01*

Mostly, the predictor variables spatial distribution, MED and ACSA_max_ impacted the model’s variance, for both low and high forces. The results confirm the influence of MED and MU spatial distribution on the number of detected MUs at low forces and are in agreement with the observations from Figures 3 and 4. ACSA_max_ showed a constant contribution at both force levels, implying that muscle size influenced the number of identified MUs.

## Discussion

In this study, we experimentally found that MED (subcutaneous fat layer), MU spatial distribution and ACSA_max_ influenced the number of MUs detected by BSS of HD-sEMG signals during low but not high force contractions. Moreover, we found a higher number of identified MUs for the ST group. This is likely due to a higher thickness of subcutaneous tissue in untrained individuals, compared to the chronically trained subjects. Moreover, the negative correlation between MED and the number of MUs is in agreement with the spatial filtering effect of the subcutaneous fat tissue, supporting previous literature [28], [43].

The subcutaneous fat increases the distance between the source (MUs) and the electrodes, resulting in more similar MUAPs shapes [9], which is a limit for the BSS, since two different sources having the same MUAPs are considered identical by the decomposition algorithm [13]. Regarding the MU spatial distribution, we observed a significant correlation between the spread in MU territories and number of MUs, at low, but not at high, contraction forces. Because the standard distance represents the spread of the MUs in space, our findings are explained by a greater difference between MUAPs when the MU territories are farther apart.

We hypothesized an effect of synchronization on the number of decomposed motor units. Although we observed that synchronization increases with force, it did not have any relation with the number of detected MUs. This confirms that BSS algorithms are robust enough not to be influenced by temporal overlapping of MUAPs [1], [19].

With respect to ACSA_max_, muscle size influenced the number of detected MUs and according to the results of our multiple regression, the effect of ACSA_max_ was the same across force levels. This result is in contrast with Farina et al. (2008) [9], which found in simulation that muscle size does not affect the discrimination of individual MUAPs. This indicates the potential limitations of simulation studies with respect to experimental results. The multiple regression model highlights the effects of MED and MU spatial distribution on the number of detected MUs, with higher significance at lower force levels. The reduced R^2^, F-statistic and p-value at high forces might indicate the influence of other unknown/unexplored parameters. Interestingly, most of our parameters were good predictors of MU numbers at low but not at high forces.

We found significant associations between different physiological parameters and the number of decomposed motor units. From all the assessed parameters, the spatial distribution of MUs, MED, and ACSA_max_ were the ones that better predicted the number of identified MUs with BSS. Our research may be useful when interpreting decomposition outcomes and when refining decomposition algorithms.

## Acknowledgments

This work was partially funded by d.hip (Digital Health Innovation Platform), a collaboration between Siemens Healthineers, Medical Valley, University Hospital Erlangen, and Friedrich-Alexander University (AdV) and by the EPSRC (project NISNEM) (DF).

